# Facility delivery and postnatal care services use among mothers who attended four or more antenatal care visits in Ethiopia: further analysis of the 2016 Demographic and Health Survey

**DOI:** 10.1101/347153

**Authors:** Gedefaw Abeje Fekadu, Fentie Ambaw Getahun, Seblewongiel Ayenalem Kidanie

**Affiliations:** School of Public Health, College of Medicine and Health Sciences, Bahir Dar University; Department of Social Work, Faculty of Social Sciences, Bahir Dar University

**Keywords:** antenatal care, facility delivery, postnatal care, quality of care, EDHS, Ethiopia

## Abstract

**Introduction:** In Ethiopia, many mothers who attend the recommended number of antenatal care visits fail to use facility delivery and postnatal care services. This study identifies factors associated with facility delivery and use of postnatal care among mothers who had four or more antenatal care visits, using data from the 2016 Ethiopian Demographic and Health Survey (EDHS).

**Methods:** To identify factors associated with facility delivery, we studied background and service-related characteristics among 2,415 mothers who attended four or more antenatal care visits for the most recent birth. In analyzing factors associated with postnatal care within 42 days after delivery, the study included 1,055 mothers who attended four or more antenatal care visits and delivered at home. We focused on women who delivered at home because women who deliver at a health facility are more likely also to receive postnatal care as well. A multivariable logistic regression model was fitted for each outcome to find significant associations between facility delivery and use of postnatal care.

**Results:** Fifty-six percent of women who had four or more antenatal care visits delivered at a health facility, while 44% delivered at home. Mothers with secondary or above level of education, urban residents, women in the richest wealth quintile, and women who were working at the time of interview had higher odds of delivering in a health facility. High birth order was associated with a lower likelihood of health facility delivery. Among women who delivered at home, only 8% received postnatal care within 42 days after delivery. Quality of antenatal care as measured by the content of care received during antenatal care visits stood out as an important factor that influences both facility delivery and postnatal care. Among mothers who attended four or more antenatal care visits and delivered at home, the content of care received during ANC visits was the only factor that showed a statistically significant association with receiving postnatal care.

**Conclusions:** The more antenatal care components a mother receives, the higher her probability of delivering at a health facility and of receiving postnatal care. The health care system needs to increase the quality of antenatal care provided to mothers because receiving more components of antenatal care is associated with increased health facility delivery and postnatal care. Further research is recommended to identify other reasons why many women do not use facility delivery and postnatal care services even after attending four or more antenatal care visits.

## 1. Introduction

### 1.1. Background

Worldwide, pregnancy-related complications are the main cause of death and disability among reproductive-age women. In 2015, about 300,000 women died from causes related to pregnancy and childbirth. Ninety-nine percent of the deaths occurred in developing countries. The majority of the maternal deaths (66%) were in sub-Saharan Africa [1, 2]. Ethiopia is the second--most populous African country, with one of the world’s highest maternal mortality ratios [3].

Obstetric hemorrhage, hypertension, abortion, sepsis, HIV, preexisting medical disorders, and other indirect causes like anemia are the main causes of maternal mortality both globally and in Ethiopia [2, 4–11]. These causes of death are preventable with proven cost-effective interventions. Antenatal care (ANC), skilled attendant at delivery, and postnatal care (PNC) are some of these effective interventions [12–18]. A review of evidence published in Lancet indicated that provision of effective care for women delivering in health facilities could prevent 113,000 maternal deaths each year [19].

Ethiopia has developed many strategies and programs to improve use of maternal health services. Family planning, ANC, facility delivery, and PNC services are provided free [20–24]. Regardless of these efforts, institutional delivery service use (26%) and PNC uptake (17%) has remained very low, while there has been improvement in use of ANC (ANC service use changed from 27% in 2000 to 62% in 2016) [25].

During antenatal care, women are encouraged to develop healthy behavior, including delivering in a health facility and using postnatal care services, in addition to other packages of interventions. Receiving antenatal care at least four times during a pregnancy increases the likelihood of receiving other maternal health services like facility delivery and postnatal care [26, 27].

Why has use of institutional delivery and of postnatal care services remained low while use of antenatal care services is relatively high in Ethiopia? The answer to this question could help health services in providing the continuum of maternal health care services. This study is designed to identify these factors, using data from the 2016 Ethiopian Demographic and Health Survey (EDHS).

### 1.2. Literature review

Antenatal, delivery, and postnatal care are part of the continuum of care for maternal and child survival [28–36]. Antenatal care can prevent maternal mortality and morbidity, directly or indirectly. Directly, ANC provides an opportunity for health care providers to detect and treat causes of maternal mortality as early as possible. Indirectly, ANC is an opportunity for health promotion to enable woman to use other services like health facility delivery and PNC. When the number of ANC visits increases, the woman is expected to acquire more knowledge and develop a better attitude about using other maternal health services [23, 37–39]. Previous studies have recommended four or more focused visits for antenatal care to improve maternal and neonatal outcomes [40, 41].

Antenatal care is thought to improve use of delivery and postnatal care services through three mechanisms. First, ANC increases the quality of information women will have about the importance of health facility delivery and postnatal care. Second, it increases women’s familiarity with medical personnel and health facilities, thus reducing the psychological costs related to seeking these services. The third mechanism is that ANC helps to create enabling or reinforcing habits to make use of these service [42].

Skilled attendance at delivery can prevent and treat life-threatening conditions that may occur at the time of delivery. Most maternal deaths occur within two days of delivery. A skilled attendant at delivery can monitor labor progress closely. This helps to prevent, detect, and manage life-threatening complications as early as possible [43–46]. Women in developing countries are more likely to get this care if they go to health institutions at the time of delivery [47, 48]. Postnatal care is another critical intervention to tackle most causes of maternal and child mortality. Postnatal care helps to detect and manage complications that may arise at the time of labor, delivery, or early after delivery. The postnatal period is the time to counsel the mother about maternal nutrition, breastfeeding, and other child care practices. During ANC, health workers can inform mothers about neonatal and maternal danger signs that may occur during the postnatal period. In addition, emotional and psychosocial support is provided to alleviate stress [23, 49–51].

Mounting evidence has indicated the importance of making the recommended number of ANC visits in increasing use of facility delivery and postnatal care. A systematic review and meta-analysis of evidence to identify the effect of antenatal care on use of health facility delivery in sub-Saharan African countries showed that mothers who attended at least four ANC visits were seven times more likely to deliver in a health facility [52]. Another study among four African countries reported a positive effect of the number of ANC consultations with use of a skilled birth attendant [42]. Studies in Colombia, Nepal, Bangladesh, Zambia, and Kenya indicated that women with a high number of prenatal visits are more likely to give birth in a health facility [53–57]. A study conducted in Debremarkos town, Northwest Ethiopia, reported that women who attended four or more ANC visits were about five times more likely to deliver in health facility compared with women who attended fewer than four visits [58].

Aside from ANC, other factors influence facility delivery. Mother’s age, educational attainment, residence, household wealth, birth order, and exposure to mass media are factors found associated with women’s use of delivery at a health facility after antenatal care [54, 56]. In a study conducted in Nepal, residence, economic status, educational status of the mother, and maternal occupation were found to be determinants of postnatal care use after antenatal care [59]. In Nigeria, service awareness, educational status, and marital status were determinants of using postnatal care services [60]. A study in Tanzania indicated that women with an intended pregnancy were more likely to use postnatal care [61].

### 1.3. Significance of the study

Regardless of efforts made by the Ethiopian government, use of facility delivery and postnatal care services is low. The proportions of women delivering in a health facility and receiving postnatal care are very low compared with the proportion attending antenatal care [25]. There is no study in Ethiopia based on nationally representative data to identify factors associated with health facility delivery and use of postnatal care services among women who attended the recommended number of ANC visits. Therefore, our study is designed for this purpose. The study will inform policymakers and programmers to improve the synergy between antenatal, facility delivery, and postnatal care by identifying factors associated with use of health facility delivery and postnatal care services among women who attended the recommended number of four or more ANC visits.

### 1.4. Research questions

This study was designed to answer the following research questions.

- What proportion of mothers who attended four or more antenatal care visits delivered in a health facility?
- What proportion of mothers who attended four or more antenatal care visits and delivered at home attended postnatal care?
- What factors are associated with mothers’ use of facility delivery and postnatal care services after completing four or more antenatal care visits?

### 1.5. Conceptual framework

This conceptual framework (Figure 1) was developed from the research literature. There are several categories of independent variables: socio-demographic characteristics of mothers (age at recent birth, educational status, marital status, religion, place of residence, wealth quintile, working status); child-related characteristics (pregnancy wantedness, birth order); and ANC-related factors (completeness of ANC components, perceived distance to facility). The two dependent variables of the study are health facility delivery after attending four or more ANC visits, and use of postnatal care services after attending four or more ANC visits and delivering at home. Analysis of direct association between the independent variables and the dependent variables was done separately for place of delivery among women with four or more ANC visits, and for postnatal care attendance among women with four or more ANC visits who delivered at home instead of in a health facility.

**Figure 1:**
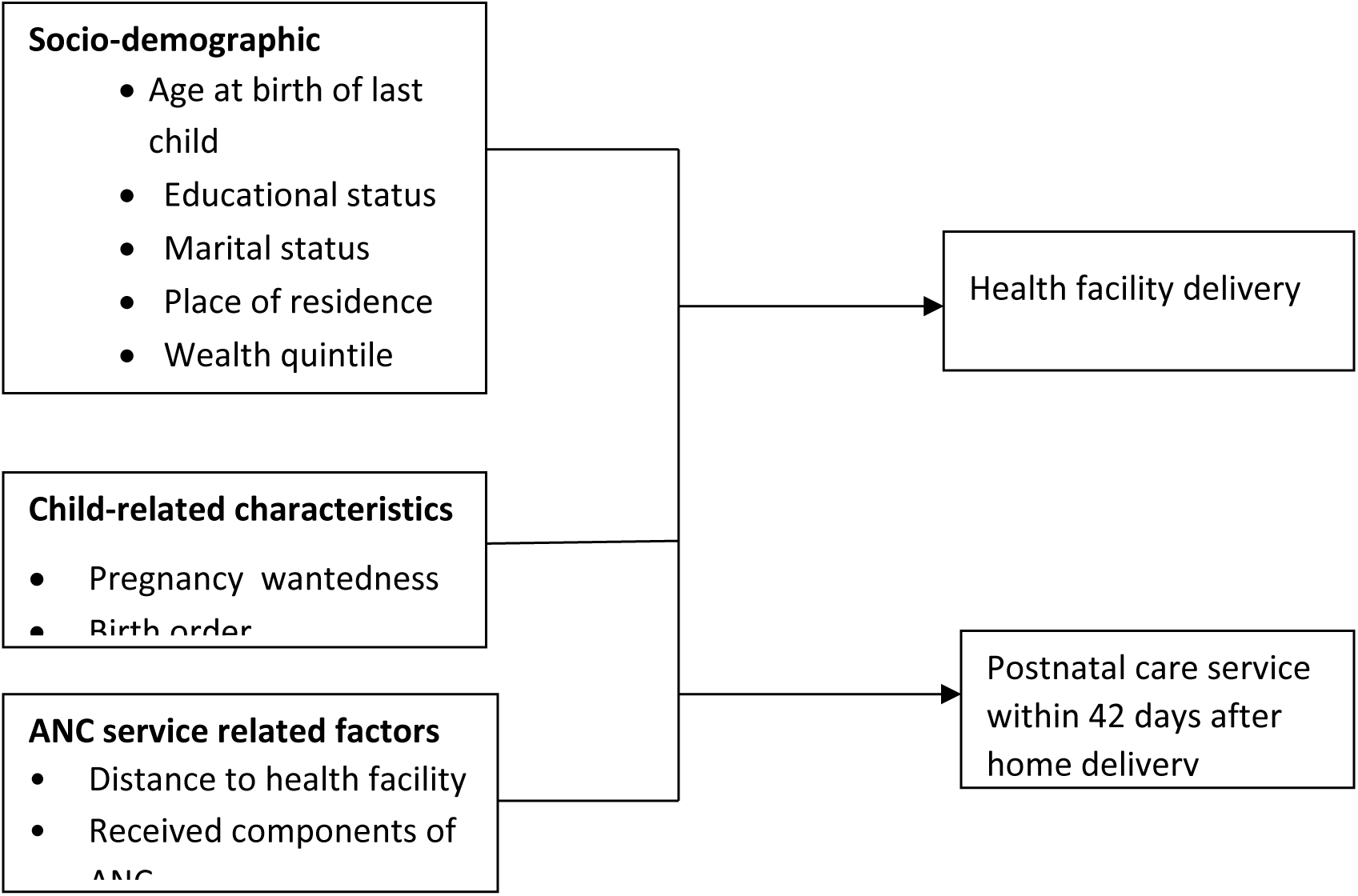
Conceptual framework of factors associated with health facility delivery and postnatal care services use among mothers who attended four or more ANC visits in Ethiopia, 2016.

## 2. Methods

### 2.1. Data

This study used data from the 2016 Ethiopian Demographic and Health Survey, which was conducted by the Central Statistical Agency (CSA), Ethiopia, and The DHS Program, ICF [62]. The 2016 EDHS was designed to provide representative data for the country as a whole and for nine regional states and two city administrations of Ethiopia. The sample design was done in two stages. In the first stage, each region was stratified into urban and rural areas and clusters were selected from both rural and urban areas based on the 2007 Ethiopian population and housing census using a probability proportional to size selection. A list of all the households was prepared in all the selected clusters. The second stage of selection used the list of households as a sampling frame and systematically selected a fixed number of households per cluster. Then, all women age 15-49 who were either permanent residents of the selected households or visitors who stayed in the household the night before the survey were included.

For this study, women who were considered for analysis were selected based on the following criteria:

- Had at least one birth in the five years before the interview
- Attended four or more antenatal care visits during the most recent pregnancy

Of the total 15,683 women included in the 2016 EDHS, 2,415 who attended four or more ANC visits for their most recent birth were included, to identify factors associated with facility delivery. From these mothers, 1,055 who delivered at home were studied to identify factors associated with receiving postnatal care (Figure 2). We focused on women who delivered at home because women who deliver at a health facility are more likely to receive postnatal care, which is initiated by health care providers and thus does not measure women’s health seeking behavior [63–66].

**Figure 2:**
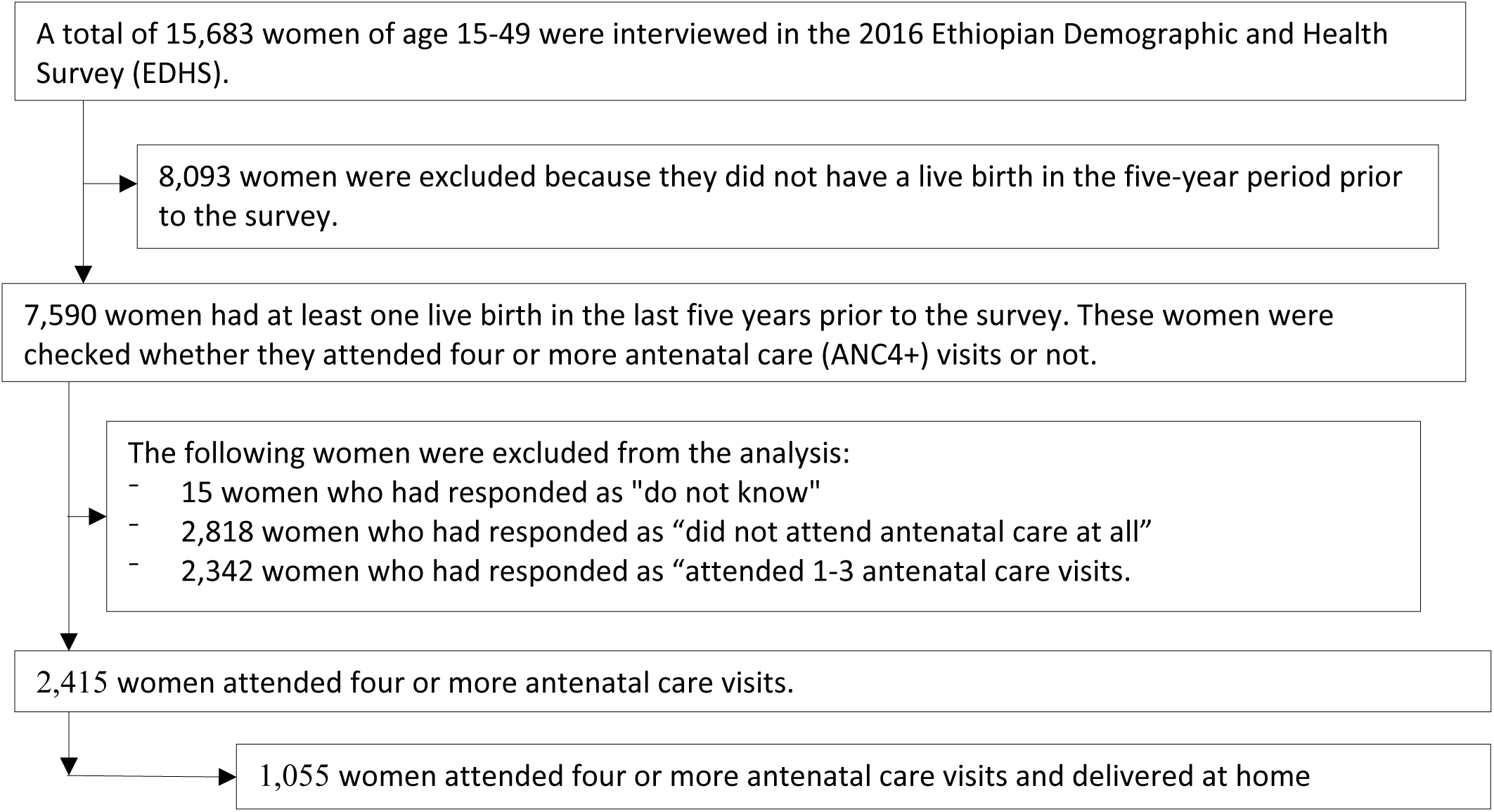
Selection of analysis sample.

### 2.2. Measurements

Outcome variables:

- Facility delivery: It was coded as “yes” if the mother gave the most recent birth at a health facility; otherwise, it was coded as “no”.
- Postnatal care within 42 days after delivery: It was coded as “yes” if the mother received PNC within 42 days after home delivery; otherwise, it was coded as “no”.

Independent variables:

- Socio-demographic characteristics: This included maternal age at the birth of the most recent child, religion, educational status, religion, marital status, place of residence (urban-rural), household wealth quintile, and working status of the mother.
- Fertility related characteristics: This included birth order of the recent child categorized as “first”, “2- 4”, or “five and above”, and wantedness of the pregnancy of the recent child categorized as “wanted then”, “wanted later”, and “wanted no more”.
- ANC service-related characteristics: This included mother’s perceived distance of the nearest health facility, categorized as “big problem” or “not a big problem”; components of antenatal care, measured by computing a summative composite score from the following antenatal components: blood pressure measurement taken (yes/no), urine sample taken (yes/no), blood sample taken (yes/no), mother told about pregnancy complications (yes/no), mother told about birth preparedness plan (yes/no), and mother received nutrition counseling (yes/no). For each of the component indicators, a woman received a point if she received the component. A total composite score was created by summing up the points for all six indicators. The total score ranges from zero to six where zero represents that the woman received none of the six antenatal care components and six represents that the woman received all six components.

### 2.3. Analytical methods

Data were analyzed using Stata 15.1. Sample weight was applied in all the analysis. Descriptive statistics were used to describe the characteristics of mothers. Multicollinearity was checked using variance inflation factors (VIF). Binary logistic regressions were used to identify significant predictors of facility delivery and postnatal care.

## 3. Results

### 3.1. Sample characteristics

A total of 2,415 mothers who had four or more antenatal care visits for the most recent birth in the five years prior to the survey were included in this analysis. As Table 1 shows, the mean age of the mothers was 27.5 years. Most of the mothers (75%) were rural residents. Almost all (93%) were either married or in union. By religion, 69% were Christian. Regarding the level of education, almost half of the mothers (48%) had no formal education, and 34% had only a primary education. The majority of the mothers (66%) had no formal job outside the home. Considering the wealth index, 30% were in the richest wealth quintile while 13% were in the poorest quintile (Table 1).

**Table 1:**
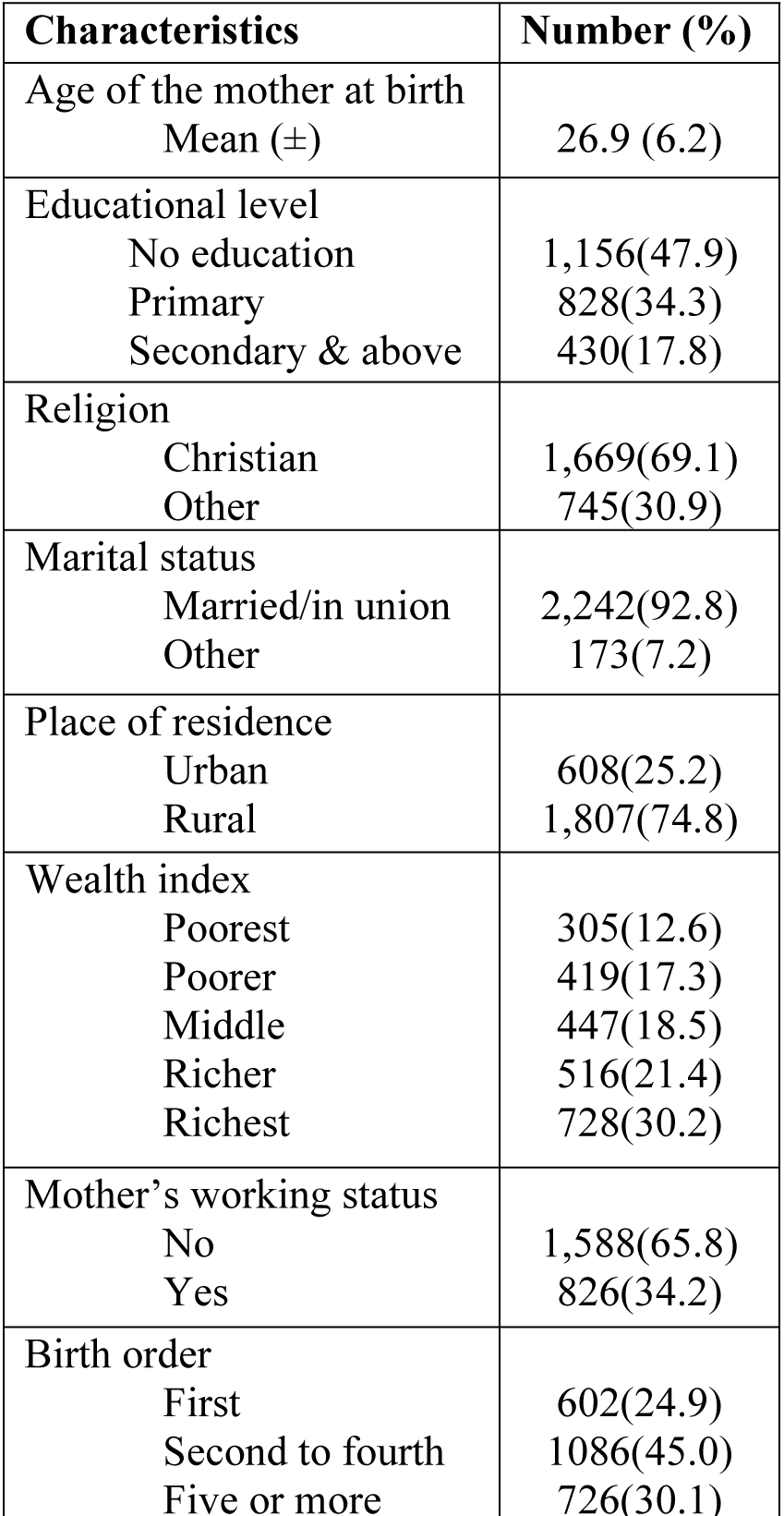
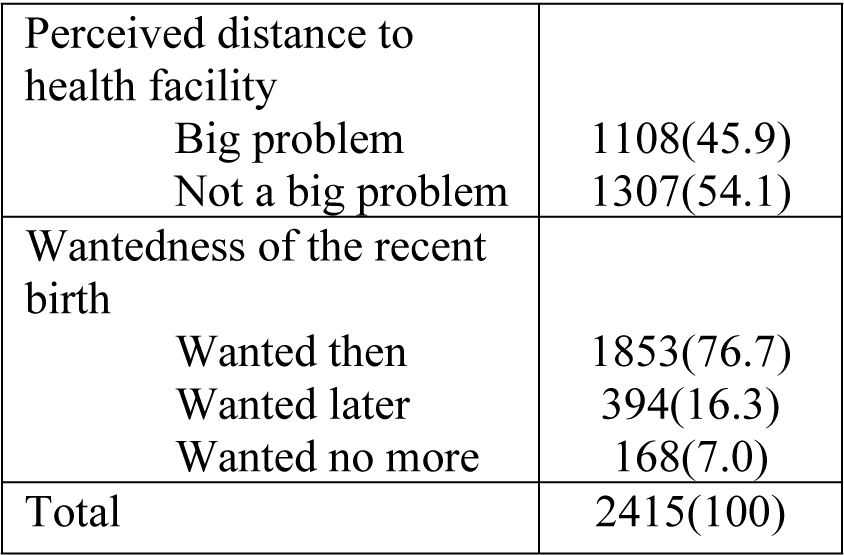
Socio-demographic characteristics of mothers who had four or more ANC visits for the most recent birth within five years before the survey, Ethiopia, 2016

For almost half (45%) of the women, the most recent birth was a second, third, or fourth birth, while for 30% it was a fifth birth or more. For 25% of the women the birth was a first birth. More than half of the mothers (54%) did not perceive distance to a health facility as a big problem for use of health services. Regarding the wanted nature of their recent pregnancy, 77% of mothers said it was wanted then, while 16% said they wanted the child later. Seven percent said the birth was unwanted.

### 3.2. Health facility delivery after four or more antenatal care visits

Among women who attended four or more ANC visits, 56% delivered at health facility, which is much higher than the level among all women (32%). Among mothers who delivered at a health facility, almost all (95%) gave birth at a government facility (not shown in the table).

As Table 2 shows, facility delivery is similar among married and non-married mothers, at just over half of women attending four or more ANC visits. Among Christian mothers, 59% delivered in a health facility. Among mothers with secondary and higher education, the great majority (91%) delivered in a health facility compared with less than half (43%) of mothers with no education. Among urban mothers, 92% had a facility delivery versus 44% of rural women. A large majority of mothers in the richest wealth quintile (88%) delivered in a health facility compared with just 34% of mothers is the poorest quintile. Among mothers who reported working at the time of interview, 70% delivered in a health facility compared with about half of non-working mothers. About two-thirds (67%) of mothers who perceived distance to a health facility as not a big problem gave birth at health facility compared with 44% of women who said distance to a health facility was a big problem (Table 2).

**Table 2:**
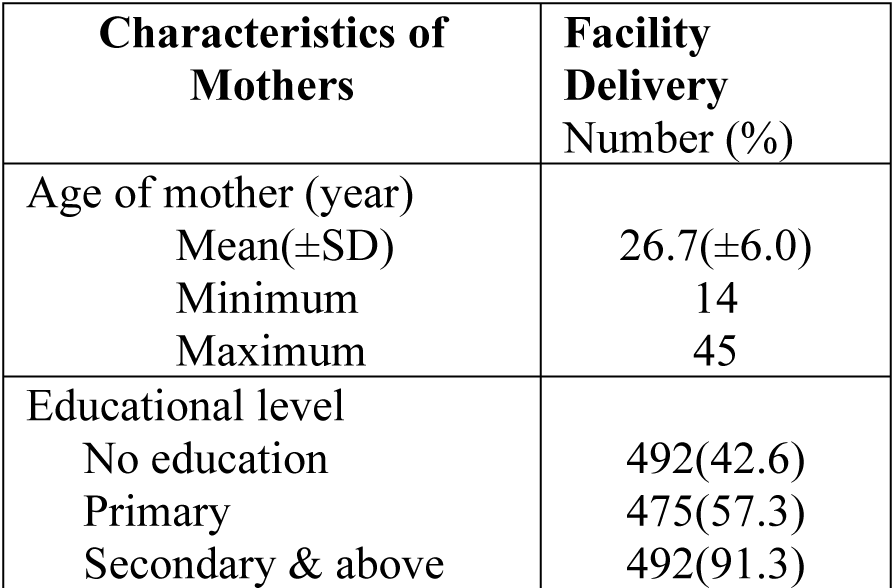
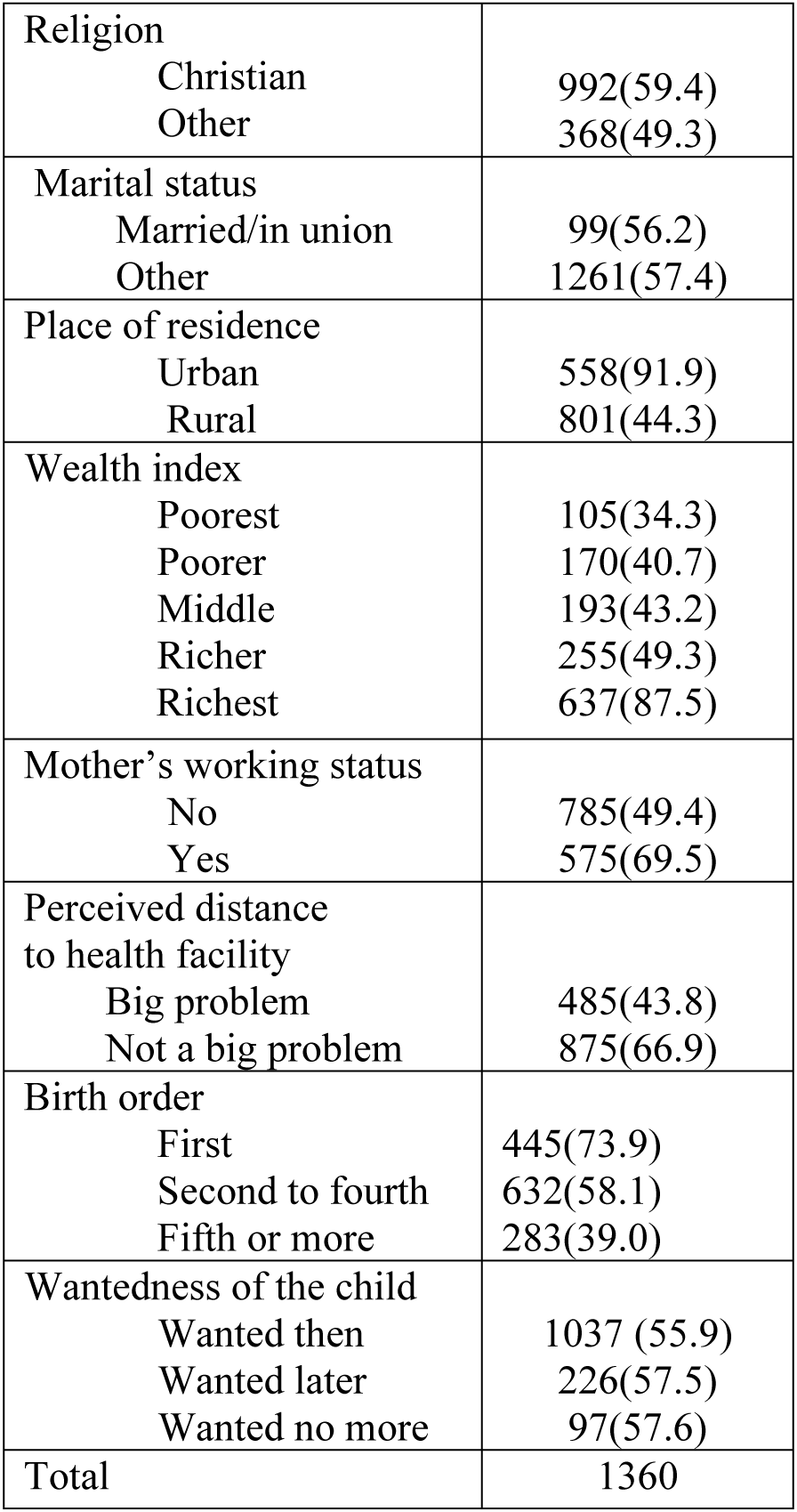
Among women who attended four or more ANC visits for the most recent birth in the five years before the survey, the percentage who delivered at a health facility by socio-demographic characteristics of mothers, Ethiopia, 2016 DHS

| Characteristics of Mothers | Facility Delivery Number (%) |
| --- | --- |
| Age of mother (year) |  |
| Mean(±SD) | 26.7(±6.0) |
| Minimum | 14 |
| Maximum | 45 |
| Educational level |  |
| No education | 492(42.6) |
| Primary | 475(57.3) |
| Secondary & above | 492(91.3) |
| Religion |  |
| Christian | 992(59.4) |
| Other | 368(49.3) |
| Marital status |  |
| Married/in union | 99(56.2) |
| Other | 1261(57.4) |
| Place of residence |  |
| Urban | 558(91.9) |
| Rural | 801(44.3) |
| Wealth index |  |
| Poorest | 105(34.3) |
| Poorer | 170(40.7) |
| Middle | 193(43.2) |
| Richer | 255(49.3) |
| Richest | 637(87.5) |
| Mother's working status |  |
| No | 785(49.4) |
| Yes | 575(69.5) |
| Perceived distance to health facility |  |
| Big problem | 485(43.8) |
| Not a big problem | 875(66.9) |
| Birth order |  |
| First | 445(73.9) |
| Second to fourth | 632(58.1) |
| Fifth or more | 283(39.0) |
| Wantedness of the child |  |
| Wanted then | 1037 (55.9) |
| Wanted later | 226(57.5) |
| Wanted no more | 97(57.6) |
| Total | 1360 |

### 3.3. Factors associated with health facility delivery after four or more antenatal care visits

During multivariable analysis, a binary logistic regression was employed. Table 3 shows that educational status, place of residence, wealth index, mother’s working status, birth order, and quality of ANC were significantly associated with facility delivery (p<0.05) among women who attended four or more ANC visits. Mothers who completed secondary and tertiary level of education had 2.9 times (95% CI: 1.6 – 5.3) the odds of health facility delivery compared with mothers without formal education. Urban residents had 3.4 times (95%CI: 1.9– 6.1) the odds of health facility delivery compared with their rural counterparts. Women in the richest wealth quintile had 2.7 times (95% CI: 1.5 – 4.8) the odds of using health facility delivery compared with mothers in the poorest quintile.

**Table 3.** Factors associated with health facility delivery among women who attended four or more ANC visits, Ethiopia, 2016.

| **Characteristics of Mothers** | **Facility Delivery**<br>**AOR(CI)** |
| --- | --- |
| Age of the mother at birth | 1.0 (1.0 – 1.1) |
| Educational level |  |
| No education | 1.0 |
| Primary | 1.1(0.8 – 1.5) |
| Secondary & above | 2.9(1.6 - 5.3)*** |
| Religion |  |
| Christian | 1.00 |
| Other | 1.1(0.8 – 1.6) |
| Marital status |  |
| Married/in union | 1.3(0.7 – 2.2) |
| Other | 1.0 |
| Place of residence |  |
| Urban | 3.4(1.9 -6.1)*** |
| Rural | 1.0 |
| Wealth index |  |
| Poorest | 1.0 |
| Poorer | 1.3 (0.8 – 2.2) |
| Middle | 1.3 (0.8 – 2.1) |
| Richer | 1.4 (0.9 – 2.2) |
| Richest | 2.7(1.5 – 4.8)*** |
| Mother's working status |  |
| No | 1.0 |
| Yes | 1.6 (1.2 -2.3)** |
| Birth order |  |
| First | 1.0 |
| 2-4 | 0.5(0.3 – 0.7)*** |
| Five or more | 0.3 (0.2 – 0.6)*** |
| Perceived distance to health facility |  |
| Big problem | 1.0 |
| Not a big problem | 1.3 (1.0 – 1.7) |
| Wanted status of child |  |
| Wanted then | 1.0 |
| Wanted later | 1.0(0.7 – 1.4) |
| Wanted no more | 1.5(0.9 – 2.5) |
| Receiving components of ANC | 1.33(1.2 – 1.4)*** |

Mothers who reported working at the time of interview had 1.6 times (95%CI: 1.2 −2.3) the odds of using facility delivery compared with non-working mothers. Women with birth order 2-4 had 50% (95%CI: 0.3% – 0.7%) lower odds delivering in a health facility compared with first-order births. Similarly, women of birth order five or more had 70% (95% CI: 0.2 - 0.6) lower odds of health facility delivery compared with first-order births. An increase in receiving components of ANC improves the odds of health facility delivery by 30% (95% CI: 1.2 - 1.4) (Table 3).

### 3.4. Characteristics of women with home delivery after four or more antenatal care visits

Among mothers who attended four or more ANC visits, 44% delivered at home. As Table 4 shows, the majority had no formal education (63%), were Christian, (64%), married or in union (93%), rural residents (95%), and had no formal job (76%). Fifty-nine percent of the women perceived that distance to health facility was a big problem to receive medical care. Among the recent births who were delivered at home, 14.9%, 43.1% and 42% were first, second to fourth, and fifth order births. The majority of recent births (77.4%) were reported wanted where as 15.9% and 6.7% were wanted latter and wanted no more respectively.

**Table 4:** Socio-demographic characteristics of mothers who delivered the most recent child at home after four more antenatal care visits, Ethiopia, 2016

| **Characteristics of Mothers** | **Home Delivery Number (%)** |
| --- | --- |
| Age of the mother in years |  |
| Mean ( ± SD) | 27.2(±6.0) |
| Minimum | 14 |
| Maximum | 46 |
| Educational level |  |
| No education | 664(62.9) |
| Primary | 354(33.5) |
| Secondary & above | 38(3.6) |
| Religion |  |
| Christian | 677(64.2) |
| Other | 378(35.8) |
| Marital status |  |
| Married/in union | 981(93.0) |
| Other | 74(7.0) |
| Place of residence |  |
| Urban | 49(4.7) |
| Rural | 1006(95.3) |
| Wealth index |  |
| Poorest | 200(19.0) |
| Poorer | 8248(23.5) |
| Middle | 254(24.1) |
| Richer | 262(24.8) |
| Richest | 91(8.6) |
| Mother's working status |  |
| No | 803(76.1) |
| Yes | 252(23.9) |
| Perceived distance to health facility |  |
| Big problem | 623(59.0) |
| Not a big problem | 432(41.0) |
| Number of ANC components received |  |
| Mean (±SD) | 3.8(±1.7) |
| Minimum | 0 |
| Maximum | 6 |
| Birth order |  |
| First | 157(14.9) |
| 2-4 | 455(43.1) |
| Five or more | 443(42.0) |
| Perceived distance to health facility |  |
| Big problem | 623(59.0) |
| Not a big problem | 432(41.0) |
| Wanted status of child |  |
| Wanted then | 817(77.4) |
| Wanted later | 167(15.9) |
| Wanted no more | 71(6.7) |
| Total | 1055 |

### 3.5. Postnatal care (PNC) among mothers who delivered at home

Among mothers who delivered at home, 2.6% (95%CI 2.2 – 3.0) received timely postnatal care versus 5.3% (95%CI 4.6-6.2) among all mothers with a recent birth. Among women who gave birth at home after attending four or more ANC visits, 8% (95% CI: 6.1% - 11.5%) attended postnatal care. The proportion who received PNC, though very small, varies across different socio-demographic characteristics. Table 5 shows that 24% of urban residents attended PNC compared with 8% of rural residents. Twelve percent of mothers who completed secondary and above education reported attending PNC versus 9% of mothers with no education. Ten percent of Christian women attended PNC versus 6% of other religions. Among the richest women, 21% attended PNC compared with 7% of the poorest women. In terms of working status, 11% of mothers working outside the home had postnatal care versus 8% of non-working women.

**Table 5:** Percentage of women who delivered at home and attended postnatal care for the most recent birth after four or more antenatal care visits by socio-demographic characteristics, Ethiopia, 2016

| **Characteristics of Mothers** | **PNC Attendance Number (%)** |
| --- | --- |
| Age of the mother in years |  |
| Mean ( ± SD) | 28.3(±6.2) |
| Minimum | 15 |
| Maximum | 43 |
| Educational level |  |
| No education | 56(8.5) |
| Primary | 28(7.9) |
| Secondary & above | 4(11.8) |
| Religion |  |
| Christian | 67(9.9) |
| Other | 22(5.7) |
| Marital status |  |
| Married/in union | 82(8.3) |
| Other | 7(9.3) |
| Place of residence |  |
| Urban | 12(24.0) |
| Rural | 77(7.6) |
| Wealth index |  |
| Poorest | 15(7.3) |
| Poorer | 8(8.63.1) |
| Middle | 28(11.1) |
| Richer | 20(7.5) |
| Richest | 19(20.5) |
| Mother's working status |  |
| No | 60(7.5) |
| Yes | 28(11.3) |
| Perceived distance to health facility |  |
| Big problem | 32(5.2) |
| Not a big problem | 56(13.1) |
| Number of ANC components received |  |
| Mean(±SD) | 5(±1.3) |
| Minimum | 0 |
| Maximum | 6 |
| Birth order |  |
| First | 8(5.2) |
| 2-4 | 45(9.9) |
| Five or more | 36(8.1) |
| Perceived distance to health facility |  |
| Big problem | 32(5.2) |
| Not a big problem | 57(13.1) |
| Wanted status of child |  |
| Wanted then | 61(7.5) |
| Wanted later | 19(11.0) |
| Wanted no more | 9(12.8) |
| Total | 89 |

**Table 6:** Factors associated with place of delivery and postnatal care after four or more ANC visits, Ethiopia, 2016.

| **Characteristics of Mothers** | **Postnatal Care Attendance**<br>**AOR(CI)** |
| --- | --- |
| Age of the mother at birth | 1.0 (1.0 – 1.1) |
| Educational level |  |
| No education | 1.0 |
| Primary | 0.9(0.4 – 1.9) |
| Secondary & above | 0. 7(0.2 – 2.9) |
| Religion |  |
| Christian | 1.0 |
| Other | 0.8(0.4 -1.7) |
| Marital status |  |
| Married/in union | 0.1.0(0.4 – 2.6) |
| Other | 1.0 |
| Place of residence |  |
| Urban | 1.3 (0.5 – 3.8) |
| Rural | 1.0 |
| Wealth index |  |
| Poorest | 1.0 |
| Poorer | 0.4(0.1 – 1.4) |
| Middle | 1.4(0.5 – 3.5) |
| Richer | 0.8 (0.3 – 2.2) |
| Richest | 1.8(0. 7 – 4.8) |
| Mother's current working Status |  |
| No | 1.0 |
| Yes | 1.4 (0.8 – 2.7) |
| Birth order |  |
| First | 1.0 |
| 2-4 | 1.5(0.4 – 5.0) |
| Five or more | 1.0 (0.2– 4. 7) |
| Perceived distance to health facility |  |
| Big problem | 1.0 |
| Not a big problem | 1.9 (1.0 – 3.8) |
| Wanted status of child |  |
| Wanted then | 1.0 |
| Wanted later | 1.30. 6 -3.1) |
| Wanted no more | 1.6(0.5 – 4.5) |
| Receiving components of ANC | 1.40 (1.1 – 1.8)* |

\*p<0.05

Considering birth order, mothers of only %, %, and %, respectively, of first, second-to-fourth, and fifthor-more births attended postnatal care. Among mothers who said their recent birth was not wanted, 13% attended PNC. Eight percent of mothers whose birth was wanted then attended PNC, as well as 11% of mothers whose birth was wanted (Tables 5).

### 3.6. Factors associated with postnatal care after ANC4^+^ visits and home delivery

From all the variables included in the multivariable logistic regression mode, only one variable showed a statistically significant association with using postnatal care services after attending antenatal care and delivering at home—receiving more components of antenatal care. An increase of one component in receiving antenatal care improves the odds of using postnatal care by 40% (95% CI: 10%-80%).

## 4. Discussion

### 4.1. Health facility delivery among women who attended four or more ANC visits

This analysis showed the level of health facility delivery among women who completed the recommended number of ANC visits. Maternal education, residence, wealth index, working status of the mother, birth order, and receiving more ANC components were important predictors of health facility delivery after four or more ANC visits.

The analysis revealed that among women who attended four or more ANC visits, just over half gave birth in a health facility. This level of use is low compared with the national plan target of 95% delivery with a skilled attendant by 2020 [67, 68]. Mothers who attended four or more ANC visits are expected to be well informed and aware of the importance of health facility delivery. They have also met the challenges of visiting a health facility. There are many possible justifications for the low level of health facility delivery among women who had the recommended number of ANC visits. The first reason may be the quality of information provided during antenatal care. The information given during ANC visits might not be comprehensive enough to attract mothers to the continuum of care. The second reason may be the women’s perceived quality of health services. If women rated the quality of ANC service low, they could be less likely to deliver in a health facility.

Mothers who completed secondary or higher level of education had higher odds of health facility delivery compared with mothers with no formal education. This finding is similar to studies conducted in Nepal, Cambodia, and in a meta-analysis in Ethiopia [54, 56, 69]. High level of education is associated with better information processing skill, improved cognitive skills and values, exerting effects on health seeking behavior through different pathways. These pathways include higher level of awareness and better knowledge of maternal health services and enhanced level of autonomy. Education also increases feelings of self-worth and confidence, which all improve service use [70, 71].

In this analysis, mothers who were urban residents had higher odds of giving birth at a health facility compared with their rural counterparts. This finding is similar to the findings of studies conducted in Ethiopia and Ghana [69, 72]. Urban women are more exposed to media messages related to the benefits of facility delivery. Cultural barriers of health service use are also less prevalent in urban areas compared with rural areas [73]. Similarity, accessibility and quality of health services may be better in urban than rural areas. Urban women are also more educated than rural woman [74–76].

The analysis revealed that women in the richest wealth quintile had higher odds of giving birth in a health facilities compared with the poorest women. This finding is consistent with other studies [54, 56, 77]. The cost of health facility delivery may not be the reason in Ethiopia, since all maternal health services are provided free. But other costs—for transport, time spent by the mother and family members at the hospital, and fear of having to make unofficial payments—could help explain the little use of health facilities for delivery among the poor. Mothers who reported working at the time of interview had higher odds of delivering at a health facility. This finding is consistent with a literature review to identify why women prefer home delivery [78]. Working mothers are more likely to be autonomous in their use of health services. They are also more likely to receive information at work that promotes health facility delivery [79–81].

Second and higher order births had lower odds of being delivered at a health facility compared with first order births. This finding is in contrast to a study in Ethiopia [82] but similar to other studies [54, 56]. One reason for this is that higher order births are more likely to be unwanted, which in turn could affect use of maternal health services [83–86]. Another reason might be that mothers could be fearful during their first birth and hence more likely to use a health facility for delivery than for subsequent births [87, 88]. In addition, women in the younger generation now initiating childbearing have more education than the older generation.

This analysis showed that mothers who receive more ANC procedures have higher odds of delivering at a health facility. This finding is similar to studies in Mexico and Cambodia [56, 89]. Another study done in Bahir Dar, Northwest Ethiopia, indicated that women who received quality ANC had higher odds of delivering at a health facility [76, 89]. The number of procedures a woman receives during ANC can be considered as a proxy indicator for the quality of ANC. During the ANC visit, the health care provider will inform the mother about birth preparedness and complication readiness plan, which implies that the woman has more information about the importance of place of delivery as a result of her ANC visit.

### 4.2. Postnatal care received among women who attended four or more ANC visits and delivered at home

Among mothers who attended four or more ANC visits, the proportion who attended postnatal care was very low compared with the targets of the national child survival and the health sector transformation plan for 2020 [67, 68]. Among women who attended four or more ANC visits and delivered at home, only 8% attended postnatal care. This implies that mothers are not using postnatal care although they managed all the challenges of attending four or more ANC visits.

Since only a small number of women attended postnatal care, many factors in the multivariable analysis were not statistically significant in their association with receipt of PNC, and others had unstable estimates. Only the number of ANC components women received during ANC visits showed a statistically significant association with postnatal care use among mothers who delivered at home after four or more ANC visits.

Mothers who received more ANC components had higher odds of also receiving PNC. Receiving more components of ANC implies that the mother is more likely to be informed about complications that may occur after delivery and thus recognize the importance of timely postnatal care.

## 5. Conclusions

The level of use of health facilities for delivery in Ethiopia among women who attended the recommended number of four or more ANC visits is low compared with the country’s national targets for 2020. Among women with four or more ANC visits, those with higher socioeconomic stats (richest, working, educated, and urban) have higher odds of delivering in a health facility. In addition, the more ANC procedures a woman receives, the more likely she is to use a health facility for delivery and to use postnatal care services.

Enabling women to complete secondary school and to work outside the home may improve health facility delivery. Economic empowerment of women may also improve health facility delivery, as the richest women had higher odds of health facility delivery in this study. Health professionals should provide more components of ANC to increase the use of institutional delivery and postnatal care. Further qualitative research is needed to identify other reasons why women who attend four or more antenatal care visits often do not continue along the continuum of care to deliver in a health facility delivery or, if they deliver at home, do not receive postnatal care.

## Acknowledgement

We are appreciative of the 2018 DHS Fellows Program facilitators, Wenjuan Wang and Shireen Assaf, who immensely assisted us on the overall development of the research project and provision of training on DHS data analysis. We would like to acknowledge the effort of the co-facilitators, Henock Yebyo and Ehab Sakr, for their support on DHS data analysis skills. We would like to extend our appreciation to the 2018 DHS fellows for their constructive feedback during the entire training sessions. We are also thankful to Bahir Dar University for allowing us to attend the DHS 2018 Fellows Program. Our acknowledgment also goes to ICF for organizing the 2018 DHS fellows workshop. We are thankful to reviewers Lindsay Mallick and Courtney Allen for their constructive comments on the paper, and to the editor, Bryant Robey, and formatter of this manuscript.

